# cSurvival: a web resource for biomarker interactions in cancer outcomes

**DOI:** 10.1101/2021.11.15.468756

**Authors:** Xuanjin Cheng, Yongxing Liu, Jiahe Wang, Yujie Chen, A. Gordon Robertson, Xuekui Zhang, Steven J. M. Jones, Stefan Taubert

## Abstract

Survival analysis is a technique to identify prognostic biomarkers and genetic vulnerabilities in cancer studies. Large-scale consortium-based projects have profiled >11,000 adult and >4,000 paediatric tumor cases with clinical outcomes and multi-omics approaches. This provides a resource for investigating molecular-level cancer etiologies using clinical correlations. Although cancers often arise from multiple genetic vulnerabilities and have deregulated gene sets (GSs), existing survival analysis protocols can report only on individual genes. Additionally, there is no systematic method to connect clinical outcomes with experimental (cell line) data. To address these gaps, we developed cSurvival (https://tau.cmmt.ubc.ca/cSurvival). cSurvival provides a user-adjustable analytical pipeline with a curated, integrated database, and offers three main advances: (a) joint analysis with two genomic predictors to identify interacting biomarkers, including new algorithms to identify optimal cutoffs for two continuous predictors; (b) survival analysis not only at the gene, but also the GS level; and (c) integration of clinical and experimental cell line studies to generate synergistic biological insights. To demonstrate these advances, we report three case studies. We confirmed findings of autophagy-dependent survival in colorectal cancers and of synergistic negative effects between high expression of *SLC7A11* and *SLC2A1* on outcomes in several cancers. We further used cSurvival to identify high expression of the Nrf2-antioxidant response element pathway as a main indicator for lung cancer prognosis and for cellular resistance to oxidative stress-inducing drugs. Together, these analyses demonstrate cSurvival’s ability to support biomarker prognosis and interaction analysis via gene- and GS-level approaches and to integrate clinical and experimental biomedical studies.

**Key points:**

- We developed cSurvival, an advanced framework using clinical correlations to study biomarker interactions in cancers, with source code and curated datasets freely available for all
- cSurvival includes new algorithms to identify optimal cutoffs for two continuous predictors to stratify patients into risk groups, enabling for the first time joint analysis with two genomic predictors;
- cSurvival allows survival analysis at the gene set (GS) level with comprehensive and up-to-date GS libraries
- The cSurvival pipeline integrates clinical outcomes and experimental cancer cell line data to generate synergistic biological insights and to mine for appropriate preclinical cell line tools
- cSurvival is built on a manually curated cancer outcomes database

## INTRODUCTION

Survival analysis, or, more broadly, time-to-event analysis, assesses the statistical association between potential risk factors and the time to an event such as death or disease recurrence [1,2]. In both basic and clinical cancer biology studies, survival analysis is an important technique for identifying prognostic biomarkers and genetic vulnerabilities. Experimentally, it is useful for hypothesis generation and mechanistic inference. Clinically, it may help stratify patients into subgroups with distinct risk profiles and guide therapeutic decision making [3,4].

Since 2006, consortium-based projects, such as The Cancer Genome Atlas (TCGA) and the Therapeutically Applicable Research to Generate Effective Treatments (TARGET), have gathered clinicopathologic data along with multi-omics molecular profiles of more than fifteen thousand adult and paediatric human tumors across diverse cancer types [5–7]. Such large data resources allow exploration into cancers at the molecular level using clinical correlations at an unprecedented scale [6].

Despite the importance of survival analysis and the richness of tumor molecular datasets, we find that currently available tools do not fully exploit the potential of survival analysis. First, existing tools can only analyze one genomic predictor at a time, typically mutation or expression of an individual gene [8–23]. However, cancer often occurs due to multiple (epi-) genomic alterations, and incorporating more than one predictor in survival analysis could identify interactions between such alterations. Such interaction analysis could also be used to screen for synthetic lethality, or to identify compensatory targets for non-targetable drivers [24–27]. This in turn could facilitate the development of combination therapies (e.g. drug cocktails), which have higher efficacy and milder side effects than mono-therapy (aka one-gene-one-drug) approaches [28,29]. Second, existing tools support analysis only at the single-gene level [8–23]; however, molecular dysregulations in cancers may involve gene sets (GSs) [30], in which a collection of genes act in concert. For example, a GS may represent a specific pathway (e.g. transforming growth factor-β (TGF-β) -mediated SMAD signaling), biological process (e.g. cell cycle), disease (e.g. hereditary nonpolyposis colorectal cancer), or treatment (e.g. chemotherapy) [31,32]. Given this, analysis of prognostic biomarkers at the GS level rather than at the single-gene level should be informative [33–35]. Third, experimentally derived cancer cell line viability data [36–38] and multi-omics profiling [39] provide valuable *in vitro* information on genetic dependencies and interactions; however, no existing tool connects clinical data to such experimental studies. This makes it difficult to identify suitable preclinical cell line tools to investigate molecular mechanisms underpinning poor prognosis.

Motivated by the lack of suitable tools to address the above challenges, we built cSurvival (Figure 1). Its major advances are:

**Figure 1.**
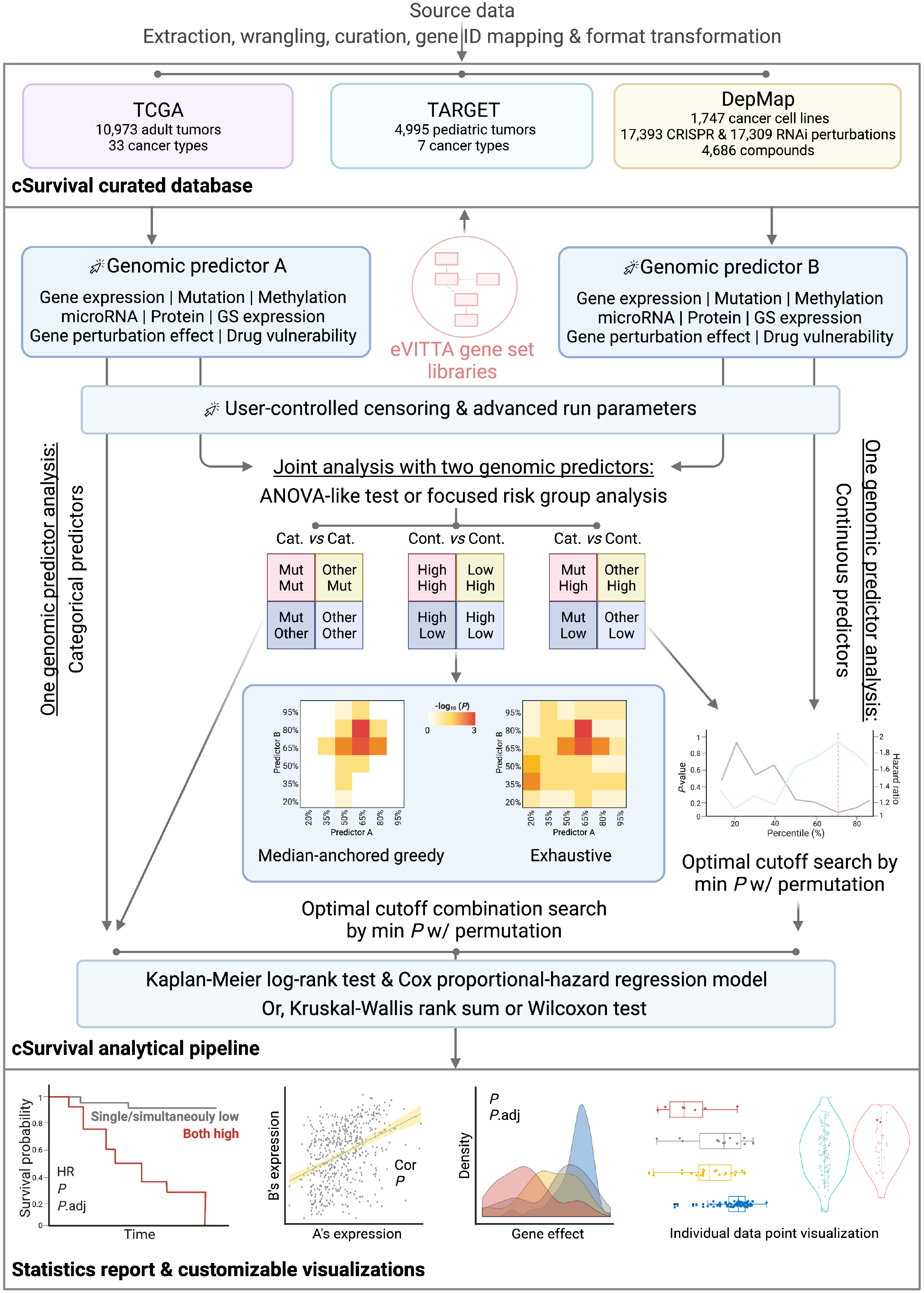
Overview of the cSurvival analytical framework. TCGA, The Cancer Genome Atlas; TARGET, Therapeutically Applicable Research to Generate Effective Treatments; DepMap, Dependency Map; GS, gene set; eVITTA, easy Visualization and Inference Toolbox for Transcriptome Analysis; Cat., categorical predictor; Cont., continuous predictor; Mut, mutated; HR, hazard ratio; *P, P*-value; *P*.adj, adjusted *P*-value; Cor, correlation coefficient.

i. Joint analysis with two genomic predictors on a wide range of individual cancer types or combinations of cancer types, including new algorithms to search for optimal cutoffs in combinations of two continuous predictors in order to stratify patients into risk groups. The two predictors can include any combination of a wide range of parameters including gene or GS expression, somatic mutation, microRNA expression, DNA methylation, and protein expression;
ii. Survival analysis at the GS level with comprehensive and up-to-date GS libraries from the easy Visualization and Inference Toolbox for Transcriptome Analysis (eVITTA) project [40]; and
iii. A pipeline to integrate clinical outcomes and experimental cancer cell line data.

We have combined a curated and integrated cancer outcomes database with a refined analytical pipeline and customizable visualizations into the cSurvival webserver so that nonprogrammers can use it. In the work described here, we demonstrate cSurvival’s capabilities with three case studies. We not only recapitulated reported cancer biomarkers and their interactions, but also identified genetic regulations consistent with published studies, demonstrating that cSurvival’s advanced pipeline facilitates cancer biomarker studies.

## MATERIAL AND METHODS

### Data extraction and processing

#### TCGA

We extracted curated clinical outcome endpoints data from the TCGA Pan-Cancer Clinical Data Resource (TCGA-CDR) [6] and multi-omics molecular data from [41]. We removed 2,614 low-quality samples (Do_not_use=True) and flagged additional 507 problematic cases based on comments in the merged_sample_quality_annotations.tsv file (https://gdc.cancer.gov/node/977). We removed these 507 cases (https://tau.cmmt.ubc.ca/cSurvival/project_data/977/flagged_cases.tsv) by default, but a user can choose to include them via the web interface. Next, we used TCGA sample type codes (https://gdc.cancer.gov/resources-tcga-users/tcga-code-tables/sample-type-codes) to extract tumor samples: for solid tumors we extracted primary solid tumor (01) samples; for acute myeloid leukemia (LAML) we extracted primary blood derived tumors (03 and 09); for skin cutaneous melanoma (SKCM) we extracted both primary solid (01) and metastatic tumors (06). Then, from [41], we used: batch-corrected, upper quartile-normalized RSEM data; merged somatic mutation calls from the Multi-Center Mutation Calling in Multiple Cancers (MC3) project [42]; purity- and ploidy-corrected, gene-level, thresholded somatic copy number (CN) data; batch-corrected, reads per million (RPM) data for expressed microRNA (miRNA) mature strands; beta values from Illumina HumanMethylation27 (HM27) and HumanMethylation450 (HM450) arrays; and batch-corrected reverse phase protein array (RPPA) data. For duplicated tumor samples, for gene expression, miRNA expression, DNA methylation, and RPPA data, we calculated the geometric means as the final readouts. We further used the annotations in [43] to map HM27 and HM450 probe IDs with chromosomal coordinates and adjacent genes.

#### TARGET

We extracted clinical and multi-omics data from the NCI Genomic Data Commons [44] with TCGAbiolinks v2.16.4 [45–47]. As above, we extracted primary tumor samples (01 for solid tumors, 03 and 09 for blood-derived). We used upper quartile normalized Fragments Per Kilobase of transcript per Million mapped reads (UQ-FPKM) data for gene expression analysis, and open-access somatic mutation calls for mutation analysis. We converted Ensembl gene IDs into HUGO symbols and Entrez IDs using org.Hs.eg.db v3.11.4 [48].

#### DepMap

We extracted cell line annotation, mutation, gene expression, CN, CRISPR-Cas9, RNA interference (RNAi), and drug sensitivity data from DepMap 21Q3 [37–39,49], normalized protein expression levels from the CCLE proteomics (TS2) database (accessed on 5/19/2021) [50], and drug information from the Drug Repurposing Hub v3/24/2020 [51].

### Calculating GS expression

After a user selects a cancer type or combinations of cancer types, we first transform the normalized gene expression counts of all samples in the selected cancer type(s) into z-scores. Then, for each sample, we computed the expression of a GS as the average expression z-score of all genes within the GS [34].

### Survival analysis

#### Censoring

A user chooses a censoring time in days, months (30.4375 days), or years (365.25 days) (default: 10 years). For a selected clinical endpoint (e.g. overall survival (OS), progression-free survival (PFS)), if the time-to-event is larger than the defined time, we set censoring status to 0 and time to the defined time; if the time-to-event is smaller than or equal to the defined time, we set censoring status to 1 and time stays unchanged.

#### Survival analysis

We apply Kaplan-Meier (KM) log-rank tests (default) and Cox proportional-hazards (PH) regression models to assess the association with prognosis, using survival v3.2.11 [52]. In joint analysis with two predictors, we use the KM log-rank test (default) or Cox PH likelihood ratio test to assess the overall significance of any difference between the four subgroup combinations of two predictors (Supplementary Figure 1). In addition, we apply Cox PH regression models to assess how two predictors jointly impact outcomes by calculating the effect sizes (hazard ratios, HRs) and significances of the two predictors and their interaction from the fitted regression model. Alternatively, users select a risk subgroup of interest, then we apply KM log-rank test (default) or Cox PH likelihood ratio test to assess the difference between the selected subgroup and the rest of the cases.

#### Determining optimal cutoffs for continuous predictors

By default, for analysis with a single continuous predictor (gene expression, miRNA expression, DNA methylation, protein expression, cell line unthresholded CN) and joint analysis with combinations of continuous and categorical predictors, we determine optimal cutoffs using the minimum *P*-value (default: KM log-rank) method [53] by testing from the lowest (default 0.2) to the highest (default 0.8) percentile with a defined step (default 0.1). In joint analysis with combinations of two continuous predictors, we determine optimal cutoffs using a median-anchored greedy (default) or an exhaustive search (described below, and in Figure 1).

Because multiple tests are conducted in searching for optimal cutoffs, we apply a *P*-value correction method to control for false positive probability (described below).

- **Median-anchored greedy search:** We construct a 2D grid using percentiles of both predictors (Figure 1). Next, we determine the starting point for greedy search by locating the minimum *P*-value computed from testing each percentile in predictor B against the median percentile in predictor A. Then, we test the nearest three unexplored points; if a lower *P*-value is found, we move the search to that newly found minimum *P*-value point and test the nearest unexplored points until no lower *P*-value can be found. We only test percentile combinations giving at least 10% (default) of total cases in each subgroup or subgroup combination.
- **Exhaustive search:** We construct a 2D grid using percentiles of both predictors (Figure 1). Next, we determine the optimal percentile combination by locating the minimum *P*-value computed from testing each percentile in predictor B against each percentile in predictor A. As above, we only test percentile combinations giving at least 10% (default) of total cases in each subgroup or subgroup combination.

#### Permutation-based multiple testing adjustment for optimally selected cutoffs

We use permutations to correct the multiple testing arising from assessing a sequence of candidate cutoffs with the minimum *P*-value method [54]. Briefly, we randomly permute outcomes values (in TCGA and TARGET, survival days and censoring status; in DepMap, gene perturbation effects or drug sensitivity scores) over the samples, and determine the new optimal cutoff. We repeat this a defined number of times (default: n = 100) to generate the null distribution of the minimum *P*-values, i.e. the empirical distribution for the minimum *P*-values when there is no association between the biomarkers and the survival outcomes. Then, we calculate an empirically adjusted *P*-value (*P*.adj) by comparing the observed minimum *P*-value to this empirical null distribution. To speed up the calculation, we use mclapply v4.0.3 [55] for parallel processing.

### Differential dependency and cell viability analysis

For two-group comparisons, we use a two-tailed two-sample Wilcoxon test (wilcox.test [55]) to assess the differences in dependency scores (CRISPR-Cas9, RNAi) or cell viabilities (drug sensitivity assays). For comparisons between more than two groups, we use a Kruskal-Wallis rank sum test (kuskal.test [55]) to test the overall significance of any difference between subgroups. For continuous genomic predictors, we determine optimal cutoffs and apply multiple testing adjustment, as described above.

### Customizable and interactive visualizations

We generate survival curves and forest plots with survminer v0.4.9 [56]. We also create interactive visualizations for further analysis with ggplot2 v3.3.5 [57] and plotly v4.9.4.1 (https://plotly.com/) (Figure 1): (a) density and box plots showing distribution of dependency scores (DepMap); (b) line plots showing *P*-values and HRs tracked over percentiles; (c) heatmaps showing *P*-values and HRs searched over percentile combinations; (d) bar plots showing distribution of somatic mutations; (e) scatter plots analyzing correlations between two continuous predictors; and (f) violin plots assessing differences in values of a continuous predictor between two categories (e.g. expression differences of a pathway between mutated *vs*. nonmutated groups). Each visualization is customizable with its own plotting parameters (e.g. colors, time intervals on the x-axis), and data points of interest are searchable and highlightable in box, scatter, and violin plots.

#### Correlation analysis in scatter plots

We use Pearson’s product-moment correlation (default), Kendall’s rank correlation tau, and Spearman’s rank correlation rho (cor.test [55]) to measure correlations between two continuous predictors.

#### Group mean analysis in violin plots

We use a two-tailed two-sample Wilcoxon test (wilcox.test [55]) to assess the differences in a continuous predictor (e.g. gene expression) between two subcategories of a categorical predictor (e.g. loss-of-function mutations *vs*. other).

### Web interface implementation

We implement the web interface of cSurvival using Apache (v2.4.29, https://httpd.apache.org), R (v4.0.3, https://www.r-project.org/), R Shiny (v1.5.0, https://CRAN.R-project.org/package=shiny), and R Shiny Server (v1.5.14.948, https://rstudio.com/products/shiny/download-server/). We use plumber (v1.1.0, https://CRAN.R-project.org/package=plumber) and pm2 (v5.1.1, https://pm2.keymetrics.io/) to host cSurvival’s API.

## RESULTS

In cSurvival v1.0.0, we have aggregated the following data. From the TCGA and the TARGET projects, clinical and multi-omics data of 10,973 adult and 4,995 paediatric tumors across 40 cancer types (33 adult, 7 paediatric). From the Dependency Map (DepMap) project [36], genetic perturbation data from 17,393 and 17,309 genes screened via CRISPR-Cas9 and RNAi in 1,032 and 712 cell lines, respectively, as well as cell viability data from 4,686 drug compounds screened in 578 cell lines and multi-omics data of 1,747 cell lines. From the eVITTA project v1.2.13 [40], 120,953 GSs (Figure 2).

**Figure 2.**
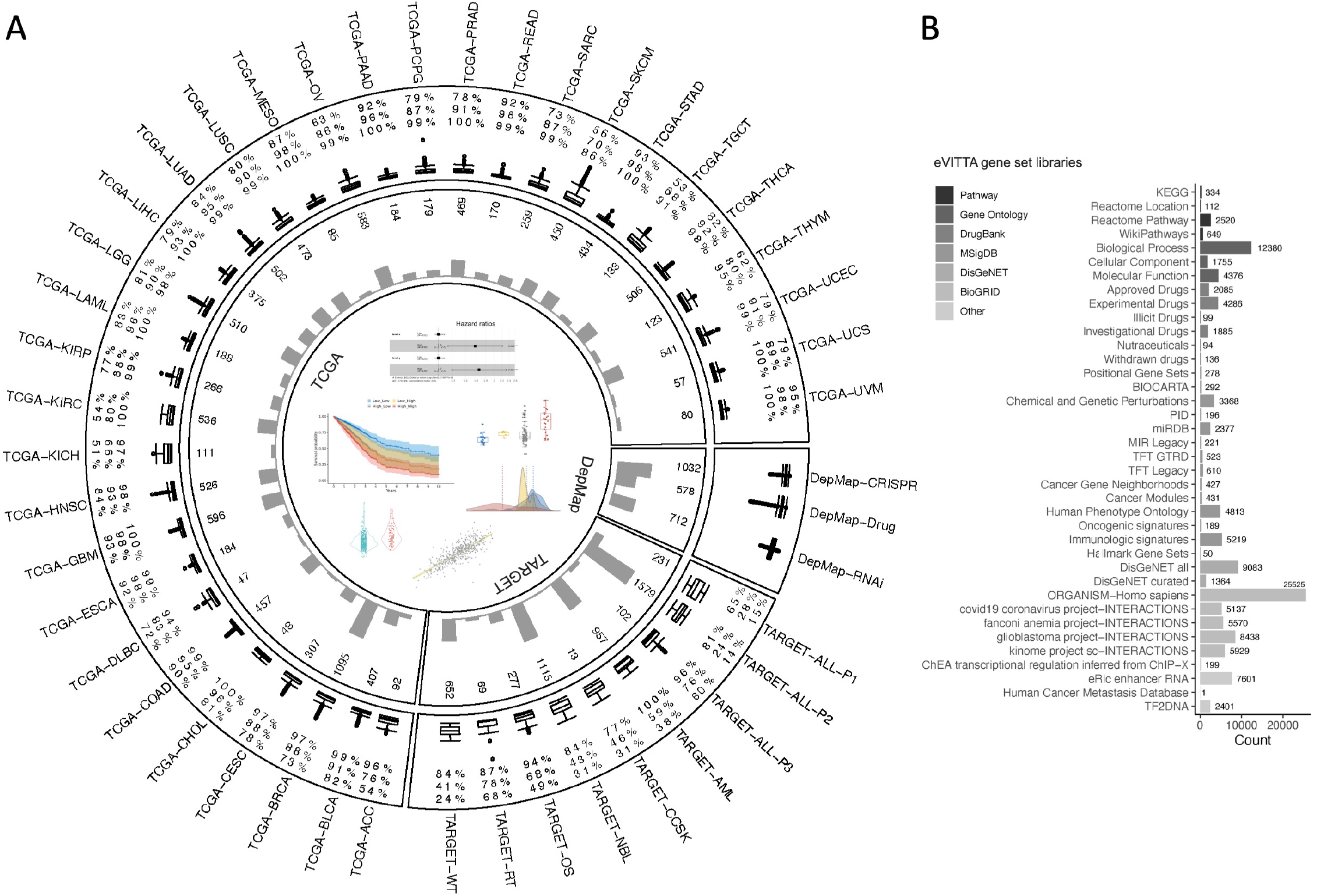
Overview of the cSurvival database. **(A)** The circos plot (rendered with circulize [74]) shows the distribution of tumor and cell line datasets: Inner to outer: example outputs; project names; histograms showing total number of cases per study; box plots showing distribution of survival days (TCGA, TARGET) or dependency scores/cell viabilities (DepMap), numbers denoting 3-, 5-, and 10-year survival rates from inner to outer; study names. **(B)** The histogram shows the distribution of GS libraries from eVITTA; numbers denote the number of GSs per library. TCGA, The Cancer Genome Atlas; TARGET, Therapeutically Applicable Research to Generate Effective Treatments; DepMap, Dependency Map; GS, gene set; eVITTA, easy Visualization and Inference Toolbox for Transcriptome Analysis.

We performed three case studies to demonstrate cSurvival’s capabilities.

First, we tested cSurvival’s analytical pipeline at the level of GSs. We recapitulated the finding that high expression of an autophagy signature (Gene Ontology [GO]: 0010506) is associated with poor overall survival in colorectal cancers (colon adenocarcinoma (TCGA-COAD) and rectum adenocarcinoma (TCGA-READ)) (Figure 3A-B, Supplementary Figure 2A) [34].

**Figure 3.**
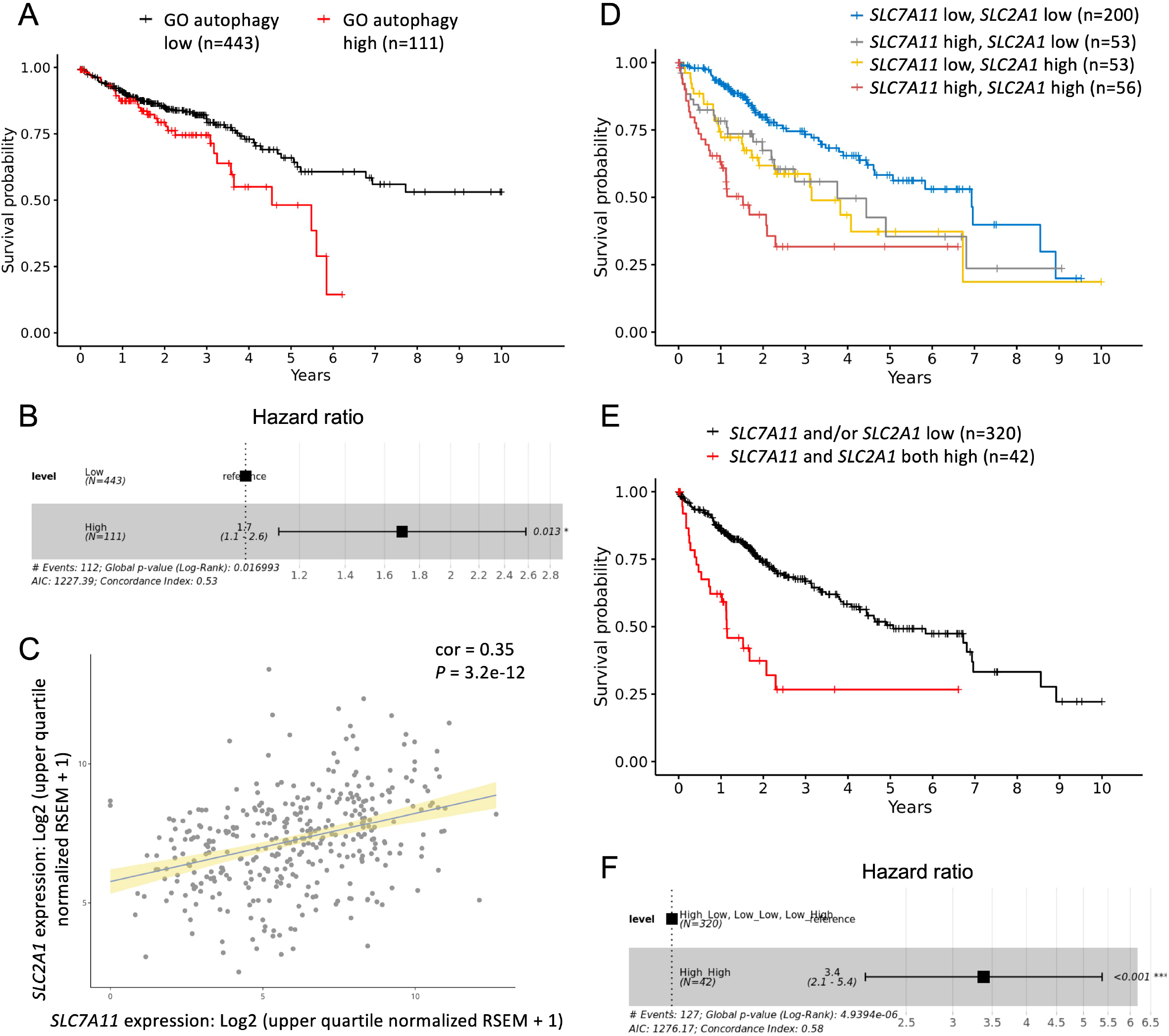
Evaluation study on autophagy-dependent survival in colorectal cancers and synergistic effects between high expression of *SLC7A11* and *SLC2A1* in liver cancers. The survival curves **(A)** and forest plot **(B)**, censored at 10 years, show correlation between GO autophagy signature (GO: 0010506) and overall survival in colorectal cancers (TCGA-COAD and READ) (*P* = 0.012, *P*.adj = 0.03, KM log-rank; HR = 1.7, *P* = 0.017, *P*.adj = 0.05, Cox PH likelihood ratio). The scatter plot **(C)** shows a moderate correlation between expression of *SLC7A11* and *SLC2A1* (Pearson’s correlation coefficient = 0.35, *P* = 3.2e-12). The survival curves **(D)**, censored at 10 years, show significant differences in overall survival rates among patients with liver hepatocellular carcinoma (TCGA-LIHC) stratified by *SLC7A11* and *SLC2A1* expression levels (*P* = 1e-07, *P*.adj < 0.01, KM log-rank; *P* = 1.7e-06, *P*.adj < 0.01, Cox PH likelihood ratio). The survival curves **(E)** and forest plots **(F)**, censored at 10 years, show a lower overall survival rate in LIHC patients with high expression of both *SLC7A11* and *SLC2A1* than in patients with low expression of *SLC7A11* and/or *SLC2A1* (*P* = 4.2e-08, *P*.adj < 0.01, KM log-rank; HR = 3.38, *P* = 4.9e-06, *P*.adj < 0.01, Cox PH likelihood ratio). GO, gene ontology; TCGA, The Cancer Genome Atlas; COAD, colon adenocarcinoma; READ, rectum adenocarcinoma; *P, P*-value; *P*.adj, adjusted *P*-value; KM, Kaplan-Meier; PH, proportional-hazard; HR, hazard ratio; LIHC, liver hepatocellular carcinoma.

Second, we performed joint analysis with gene expression data of solute carrier family 7 member 11 (*SLC7A11*) and solute carrier family 2 member 1 (*SLC2A1*, also known as glucose transporter 1 (*GLUT1*)) in liver hepatocellular carcinoma (TCGA-LIHC). We found that *SLC7A11* and *SLC2A1* showed a moderate correlation in their expressions (Figure 3C), and that patients with higher expression of both *SLC7A11* and *SLC2A1* showed significantly lower survival rates than patients with low expression of *SLC7A11* and/or *SLC2A1* (Figure 3D-F, Supplementary Figure 2B-C). The synergistic negative effects between high expressions of *SLC7A11* and *SLC2A1* on outcomes were also observed in several other cancer types (Supplementary Figure 3). These results are consistent with the finding that co-targeting the L-cystine importer *SLC7A11* and the glucose transporter *SLC2A1* induces synthetic lethal cell death in glucose-deprived cell lines [58].

Third, to illustrate cSurvival’s integrated workflow, and to show how cSurvival can bridge clinical and cell line studies, we assessed the Nrf2 (nuclear factor erythroid 2 related factor 2 (*NFE2L2*))-Keap1 (Kelch-like erythroid cell-derived protein with CNC homology-associated protein 1 (*KEAP1*)) signalling pathway. Nrf2 is a master orchestrator of oxidative homeostasis and is primarily regulated by Keap1 [59]. In cancers, *KEAP1* is frequently mutated, resulting in constitutively active Nrf2 that protects cancer cells from chemotherapeutic agents and facilitates cancer progression [59]. For example, Nrf2 is aberrantly activated in ∼30% of human lung cancers [60]. Using cSurvival, we found that expression or mutation of *NFE2L2* and *KEAP1* themselves showed no association with patient overall survival (Supplementary Figure 4). However, high expression of genes in the Nrf2-antioxidant response element (ARE) pathway (WikiPathways: WP4357) correlated strongly with poor prognosis in lung adenocarcinoma patients (Figure 4A-B). Consistent with this clinical finding, in DepMap’s experimental genetic perturbation screens, *KEAP1*-mutated lung cancer cell lines were more sensitive to *NFE2L2* knockout and knockdown (Figure 4C-F), consistent with *KEAP1* mutation being the main driver for oncogenic Nrf2 activation [61,62]. Likewise, cell lines with higher expression of Nrf2-ARE pathway genes showed greater resistance to a potent oxidative stress inducer, menadione (BRD-K78126613-001-28-5) (Figure 5A-B) [63]. Moreover, cSurvival analysis showed that the male A549 and the female H2172 cell lines both harbor deactivating *KEAP1* mutations, manifest high Nrf2-ARE pathway expression, and show relatively high resistance to menadione (Figure 5). These findings are consistent with published reports that the A549 cell line is an excellent tool for studies on Nrf2 regulation and activity [64–66]; extending this, cSurvival analysis suggests that the female H2172 cell line is another excellent model to study Nrf2 function and can be used in conjunction with A549 to study sex-specific effects (Figure 5).

**Figure 4.**
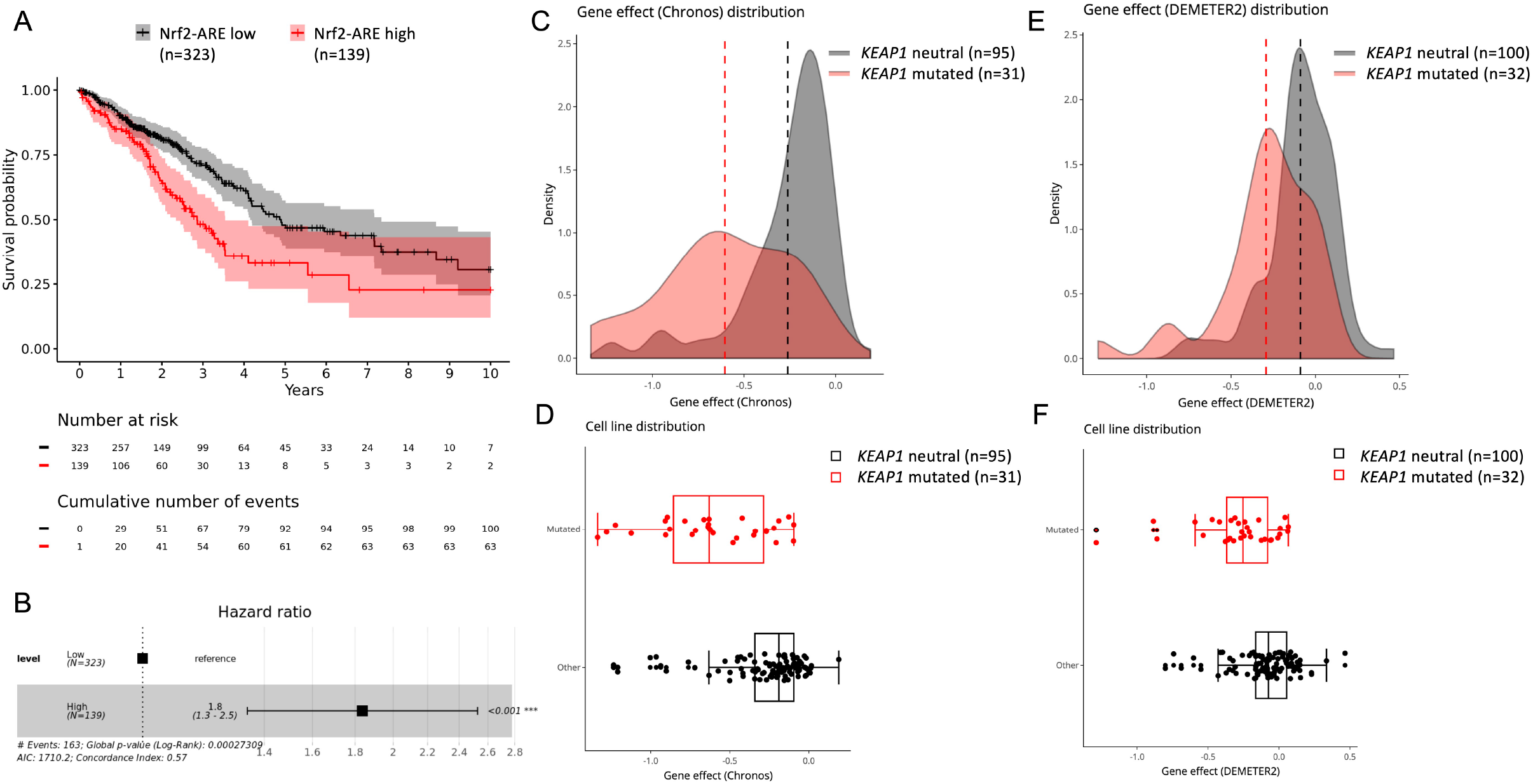
High expression of the Nrf2-ARE pathway correlates with poor prognosis and *KEAP1* mutation in lung cancers. The survival curves **(A)** and forest plot **(B)**, censored at 10 years, show correlation between high expression of the Nrf2-ARE pathway and poor prognosis in TCGA-LUAD cases (*P* = 0.00015, *P*.adj < 0.01, KM log-rank; HR = 1.84, *P* = 0.00027, *P*.adj < 0.01, Cox PH likelihood ratio). Shades reflect 95% confidence intervals in survival curves in (A). The density and box plots show lung cancer cell lines with an inactivating *KEAP1* mutation (red) being more sensitive to *NFE2L2* CRISPR-Cas9 knockout (*P* = 4.1e-07, two-tailed Wilcoxon test) **(C, D)** and RNAi knockdown (*P* = 8.4e-05, two-tailed Wilcoxon test) **(E, F)**. ARE, antioxidant response element; TCGA, The Cancer Genome Atlas; LUAD, lung adenocarcinoma; *P, P*-value; *P*.adj, adjusted *P*-value; KM, Kaplan-Meier; HR, hazard ratio; PH, proportional-hazard; RNAi, RNA interference; Chronos, an algorithm for inferring gene knockout fitness effects; DEMETER2, gene dependency estimates for RNAi datasets.

**Figure 5.**
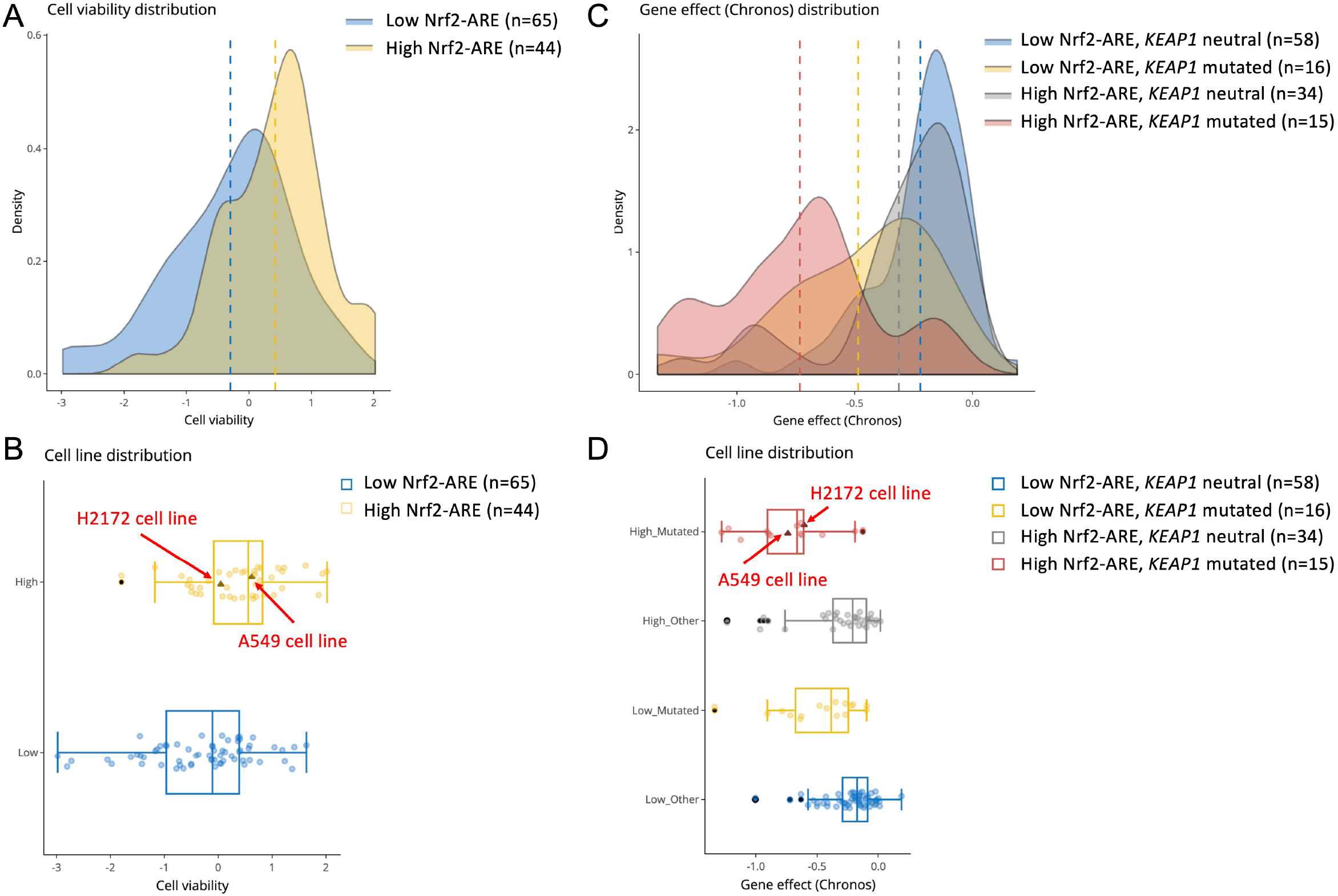
The male A549 and the female H2172 cell lines are top candidates for studies on Nrf2 regulation and activity. The density **(A)** and box **(B)** plots show that lung cancer cell lines with higher expression levels of Nrf2-ARE pathway genes exhibit higher resistance to the oxidative stress inducer menadione (BRD-K78126613-001-28-5, *P* = 0.00014, *P*.adj < 0.01, two-tailed Wilcoxon test). The density **(C)** and box **(D)** plots show that lung cancer cell lines with different *KEAP1* mutation status and expression levels of Nrf2-ARE pathway genes show sensitivity differences to *NFE2L2* knockout (*P* = 2.6e-06, *P*.adj < 0.01, Kruskal-Wallis rank sum test). The A549 and the H2172 cell lines are highlighted in triangle shape with darker color and labelled in the box plots. ARE, antioxidant response element; *P, P*-value; *P*.adj, adjusted *P*-value; Chronos, an algorithm for inferring gene knockout fitness effects.

Together, these analyses demonstrate cSurvival’s unique ability to support biomarker prognosis and interaction analysis via gene- and GS-level approaches, and to facilitate integrating clinical and experimental biomedical studies.

## DISCUSSION

Cancer arises from accumulated genetic and epigenetic alterations, creating interactions that endow cancer cells with growth and survival advantages. Such functional relationships happen not only between genes, but also between GSs, or between genes and GSs. Correspondingly, regimens that combine multiple drugs targeting different genes/GSs have emerged as more effective and less toxic than mono-therapy approaches [28,29]. Pinpointing genetic interactions in cancers is thus an important research goal.

Here, we developed cSurvival, an open-source framework to identify potential genetic interactions and survey preclinical cell line tools. cSurvival offers innovative algorithms that can assess interactions between many types of cancer biomarkers, including GSs, as well as a curated database combining clinical and experimental data to generate synergistic biological insights. As shown by the three case studies, cSurvival sheds lights on genetic interactions in cancers and facilitates identification of preclinical cell line tools for mechanistic studies.

For long-term sustainability, we have automated data extraction from the NCI Genomic Data Commons (https://gdc.cancer.gov) [44] and will continuously follow consortium efforts, so that cSurvival uses up-to-date processing pipelines, human reference genome, and gene annotations. We also plan to expand the cSurvival database, incorporating resources such as the International Cancer Genome Consortium (ICGC) [67] and the Connectivity Map (CMap) [68,69]. Future iterations of cSurvival may also address the challenges of immune [70] and ethnicity [71] heterogeneities in human populations.

Although we built cSurvival to analyze molecular biomarkers in cancers, its analytical approaches can also be applied to other types of (bio) markers and/or other diseases. For instance, its algorithms to search for optimal cutoffs for interaction analysis could also apply to data from biomedical imaging [72] or drug cocktail effect assessment [73], which sometimes involve combinations of continuous and/or categorical variables.

In summary, cSurvival offers a curated database and innovative analytical pipelines to examine cancer biomarkers at high resolution. It complements existing resources such as cBioPortal [9], and enhances mechanistic investigation of malignancy etiologies using clinical correlations. Its intuitive yet flexible web interface makes it a handy tool for experimental and clinical researchers alike.

## AVAILABILITY

cSurvival (https://tau.cmmt.ubc.ca/cSurvival) is free and open to all users and has no login requirement. The source code for building cSurvival’s framework and database is available at GitHub (https://github.com/easygsea/cSurvival.git).

## SUPPLEMENTARY DATA

Supplementary figures are available at Briefings in Bioinformatics online.

## ACKNOWLEDGEMENT

We thank Dr. P.W. Laird (Van Andel Institute, Grand Rapids, MI), T. Lichtenberg (University of Chicago, Chicago, IL), Dr. C.A. Maxwell (The University of British Columbia, Vancouver, BC), Dr. H. Shen (Van Andel Institute, Grand Rapids, MI), Dr. Z. Zhou (The Chinese University of Hong Kong, Shatin, HK), and Taubert lab members for critical comments on the manuscript. Figure 1 was created with BioRender.com, Toronto, Canada.

## FUNDING

This work was funded by grant support from The Canadian Institutes of Health Research (CIHR; PJT-153199); and the Natural Sciences and Engineering Research Council of Canada (NSERC; RGPIN-2018-05133 to S.T., RGPIN-2017-04722 to X.Z.), and a Canada Research Chair (No. 950231363 to X.Z.). S.J.M.J. acknowledges funding from the Canada Research Chairs program. Funding for open access charges: Canadian Institutes of Health Research (CIHR; PJT-153199).

**Xuanjin Cheng** is a research associate in Medical Genetics at the University of British Columbia. Her research combines bioinformatics and molecular biology to study gene regulation in development and disease.

**Yongxing Liu** is a fourth-year Mathematics student at the University of British Columbia. He is interested in data science, machine learning, and bioinformatics.

**Jiahe Wang** is currently a Computer Science Master student at Simon Fraser University with a concentration on Big Data and Machine Learning.

**Yujie Chen** is an undergraduate student majoring in Statistics at the University of British Columbia, and her research interest is Machine Learning and Bioinformatics.

**Andrew Gordon Robertson** is an analyst with BC Cancer’s Canada’s Michael Smith Genome Sciences Centre in Vancouver Canada. He has contributed to many TCGA, PanCancer Atlas, and GDAN projects and publications.

**Xuekui Zhang** is a Tier 2 Canada Research Chair in biostatistics and bioinformatics and an assistant professor in the Department of Mathematics and Statistics at the University of Victoria.

**Steven J.M. Jones** is co-director and head of bioinformatics at the Genome Sciences Centre, BC Cancer. He is also a Professor in Medical Genetics at the University of British Columbia.

**Stefan Taubert** is an associate professor in in Medical Genetics at the University of British Columbia who studies gene regulation in health and disease.

